# Cue-specific neuronal ensembles span intermittent rate coding of working memory

**DOI:** 10.1101/2023.10.06.561121

**Authors:** Matthew F Panichello, Donatas Jonikaitis, Jin Oh, Shude Zhu, Ethan B Trepka, Tirin Moore

**Author notes:** Equal Contribution (Matthew F Panichello, Donatas Jonikaitis).

## Abstract

Persistent, memorandum-specific neuronal spiking activity has long been hypothesized to underlie working memory. However, emerging evidence suggests a possible role for ‘activity-silent’ synaptic mechanisms. This issue remains controversial because evidence for either view has largely depended on datasets that fail to capture single-trial population dynamics or on indirect measures of neuronal spiking. We addressed this by examining the dynamics of mnemonic information on single trials obtained from large, local populations of prefrontal neurons recorded simultaneously in monkeys performing a working memory task. We show that mnemonic information does not persist in the spiking activity of prefrontal neurons, but instead alternates between ‘On’ and ‘Off’ periods during memory delays. At the level of single neurons, Off periods are driven by a coordinated loss of selectivity for memoranda and a return of firing rates to baseline levels. Further exploiting the large-scale recordings, we asked whether the functional connectivity among large neuronal ensembles depended on information held in working memory. We show that mnemonic information is available in the pattern of ensemble connectivity during the memory delay in both On and Off periods of neuronal activity. Intermittent epochs of memoranda-specific spiking therefore coexist with activity-silent mechanisms to span memory delays.

## Introduction

Working memory allows us to retain and manipulate information on short timescales and it is central to complex cognitive processing and adaptive behavior (Baddeley, 2000; Ehrlich & Murray, 2022). Foundational work in the 1970s revealed that working memory is associated with sustained neuronal spiking activity in primate prefrontal cortex (Fuster & Alexander, 1971). Subsequent studies demonstrated that the persistent spiking of many neurons is specific to a remembered cue (Funahashi et al., 1989; Fuster, 1973; Wimmer et al., 2014). Persistent activity has been observed during both spatial and feature-based working memory tasks (Armstrong et al., 2009; S. C. Rao et al., 1997; Wilson et al., 1993), as well as within many cortical and subcortical brain structures (Chelazzi et al., 1993; Glimcher & Sparks, 1992; Hikosaka et al., 1993; Miyashita & Chang, 1988; Snyder et al., 1997; van Kerkoerle et al., 2017). In addition to nonhuman primates, it has also been observed in multiple animal models (Inagaki et al., 2017) as well as in humans (Harrison & Tong, 2009; Vogel & Machizawa, 2004). Combined, this evidence has established persistent spiking as the dominant model of working memory (Wang, 2021).

In spite of the predominance of the persistent spiking model of working memory, an alternative class of models has received increased attention in recent years. This class of models proposes that, rather than persistent activity, working memory is instead supported by ‘activity-silent’, synaptic mechanisms (Lundqvist et al., 2011; Mongillo et al., 2008; Stokes, 2015). Specifically, information held in working memory is stored by the pattern short-term plastic changes initiated by a particular memory cue. Proof-of-principle simulations demonstrate that short-term plasticity (STP) can maintain information in the absence of persistent spiking (Lundqvist et al., 2011; Mongillo et al., 2008). Evidence of such latent memory traces has been reported using a variety of methods (Barbosa et al., 2020; Lewis-Peacock et al., 2012; Rose et al., 2016; Sprague et al., 2016; Wolff et al., 2017). For example, STP, as inferred from functional connectivity, has been shown to account for inter-trial effects and the maintenance of tasks sets during working memory (Barbosa et al., 2020; Fujisawa et al., 2008). However, evidence of cue-specific synaptic effects during canonical working memory delays has yet to be reported.

Nevertheless, synaptic models of working memory can potentially address key shortcomings of the persistent spiking model. For one, persistent activity has been reported to be modest, or even absent in some cases (Rose et al., 2016; Shafi et al., 2007; Sprague et al., 2016; Wolff et al., 2017), and to vary with task demands (Lewis-Peacock et al., 2012; Watanabe & Funahashi, 2014). Second, and more importantly, delay-period activity can be highly variable on single trials (Compte et al., 2003), prompting some to question the utility of persistent spiking as a reliable mechanism for memory maintenance (Lundqvist, Herman, & Miller, 2018; Mongillo et al., 2008; Stokes, 2015). In addition, the high-gamma component of prefrontal local field potentials appears bursty, rather than persistent, during memory delays, suggesting that population spiking may be similarly irregular (Lundqvist et al., 2016; Lundqvist, Herman, Warden, et al., 2018). In principle, a synaptic mechanism could eliminate, or at least minimize, disruptions in memory maintenance due to spiking irregularities. However, the relative contributions of spiking and synaptic mechanisms to working memory remain largely unresolved.

To address the above questions, we studied the activity of neurons within dorsolateral prefrontal cortex in two monkeys (A and H) (Supplementary Materials). Monkeys were trained to perform two variants of a spatial working memory task (Fig. 1a). In both variants, the monkey was first presented with a brief (50 ms) spatial cue at one of eight possible locations while fixating a central spot. Following the cue, the monkey maintained fixation during a memory delay (1400-1600 ms). In one task (match-to-sample)(Hasegawa et al., 2004), two targets appeared after the delay and the monkey was rewarded for making an eye movement to the target appearing at the previously cued location. In the second task (memory-guided saccade)(Funahashi et al., 1989), no targets appeared after the delay and the monkey was rewarded for making an eye movement to the previously cued (blank) location. Both monkeys successfully learned the two tasks (Fig 1b). Trials of both task types were randomly interleaved and were pooled for subsequent analyses.

**Figure 1.**
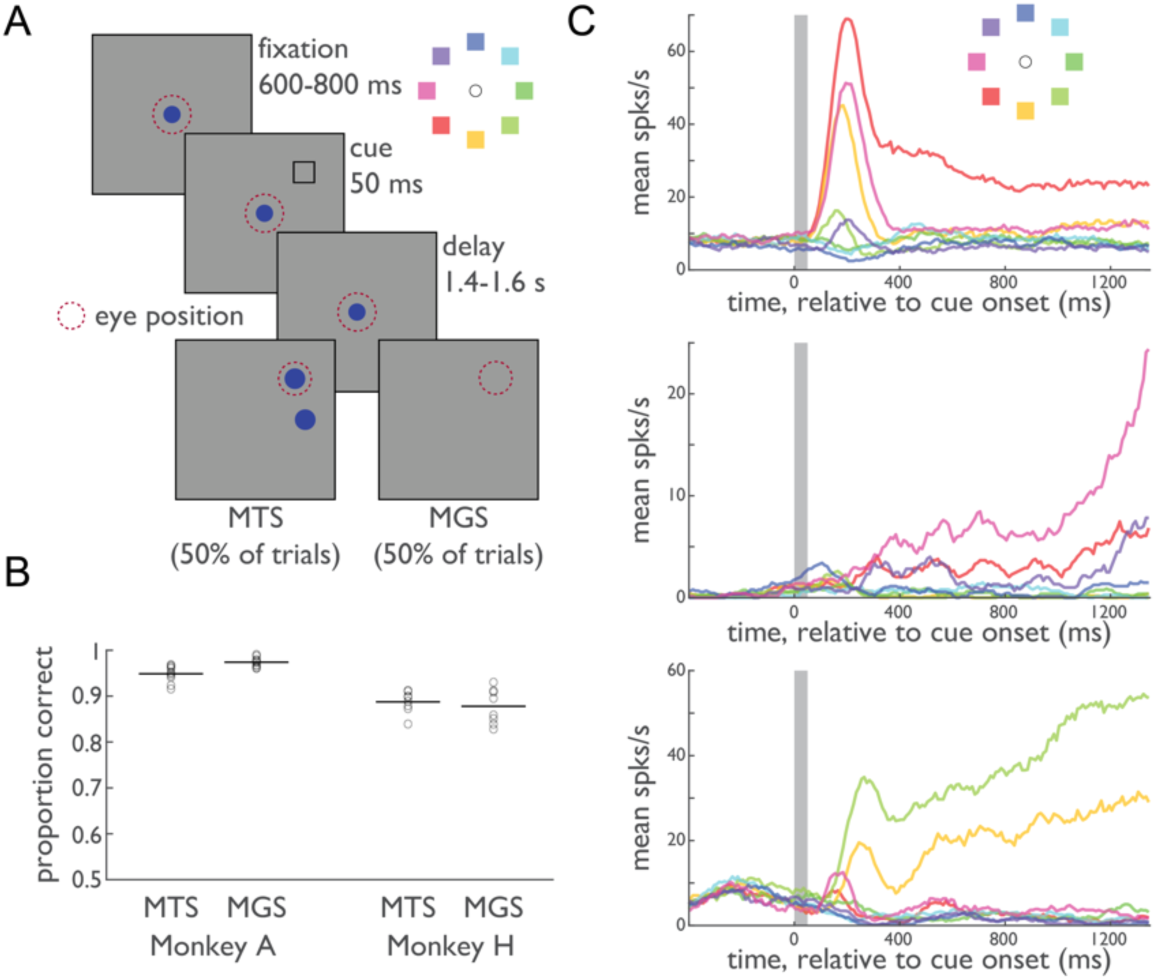
Persistent, trial-averaged neuronal responses during spatial working memory. (A) Delayed match-to-sample (MTS) and memory-guided saccade (MGS) tasks. On each trial, the animal was presented with a cue at one of eight possible locations (inset). After a memory delay period, the animal received fluid reward for making an eye movement to the previously cued location. (B) Proportion correct for the MTS and MGS tasks. Circles denote individual sessions; lines show mean across sessions. (C) Trial-averaged peristimulus time histograms for 3 example prefrontal neurons displaying canonical persistent activity during the memory delay period. Colors denote the different cue locations (inset).

### Large-scale recordings from local populations of primate prefrontal neurons

As expected from previous studies (Funahashi et al., 1989; Fuster, 1973), we observed a substantial proportion of prefrontal neurons with cue-specific memory delay activity (mean 49% per session). Trial-averaged responses of these neurons suggest that their firing rates are sufficient to encode the remembered cue during the memory delay (Fig. 1c). However, trial-averaging can obscure the high variability of spiking activity present on single trials (Lundqvist, Herman, & Miller, 2018). Thus, for individual neurons, cue information may be unreliable at times during the delay. Nonetheless, it could be that lapses in cue information for some neurons in the population are compensated by the continued activity of other neurons encoding the same memorandum. Alternatively, these lapses could be coordinated such that cue information fails to persist throughout the memory delay across the entire neuronal population. To distinguish between these possibilities, it is crucial to simultaneously measure the activity of large populations of neurons and examine their activity on single trials in order to evaluate the contribution of persistent spiking to working memory.

In the past several years, high-density, silicon probes, most notably Neuropixels probes (IMEC inc.), have revolutionized large-scale electrophysiological recordings in the mouse brain (Jun et al., 2017; Steinmetz et al., 2021). More recently, these probes were adapted for use in nonhuman primates (Trautmann et al., 2023). We used these newly developed probes to obtain recordings from large, dense populations of prefrontal neurons in monkeys performing the spatial working memory tasks (Fig. 2A). Our Neuropixels recordings (N=18 sessions) typically yielded 100s of single and multi-units in each session (mean = 272 +/-53; N = 4,894 total) (Supplementary Materials). In addition to memory delay neurons, these recordings allowed us to capture the spatial distribution of multiple functional classes of neurons (Fig. S1). For example, neurons selective to multiple task components (e.g., visuomotor neurons) tended to be more closely spaced than neurons selective to only one.

**Figure 2.**
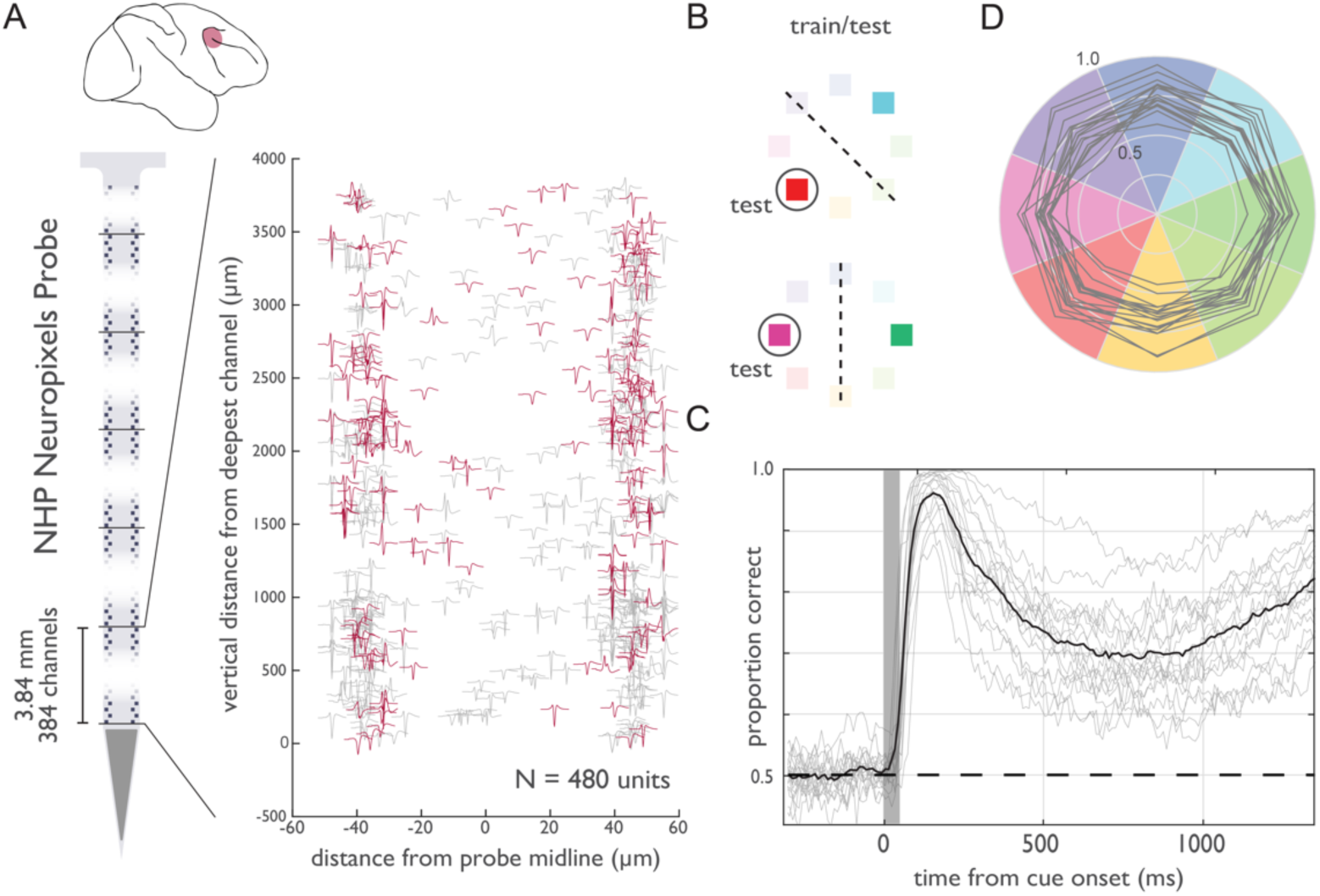
High-density neuronal recordings from prefrontal cortex. (A) Top, location of recordings in lateral prefrontal cortex (inset). Left: Schematic of the Neuropixels NHP probe, highlighting the contiguous block of 384 active channels near the probe tip. Right: spike waveform templates for 480 single- and multi-units extracted from a single example recording session in Monkey A, shown at their measured location on the probe surface. Units plotted in red showed selectivity for cue location during the delay period. (B) Leave-one-trial-out training procedure. For each trial and time point, a classifier was trained on the remainder of trials to discriminate the same cue location as the test trial from the opposite cue location. (C) Mean proportion correct, averaged across the memory period (+500 to +1400 ms relative to cue onset) by cue locations. Gray traces show individual sessions. (D) Classification accuracy (proportion of trials correct) for cue location, relative to cue onset. Gray traces indicate individual sessions.

Most importantly, these recordings allowed us to quantify the information that local populations of neurons collectively conveyed about the remembered cue location. To do this, we used a leave-one-out, binary classification procedure. For each trial within a session, we trained logistic regression models to discriminate the test location from its opposite location across the trial duration (Fig. 2B). For these and subsequent analyses, the ‘memory period’ was defined as the period from 500 to 1400 ms after the cue appearance (Supplementary Materials). Across recording sessions, mean classification accuracy was significantly above chance throughout the memory period (all p < 0.001, sign-rank)(Fig. 2C). Moreover, for each individual session, mean classification across the memory period exceeded chance performance (all p <0.001, sign rank), with accuracies ranging from 61% to 89%. Lastly, classification accuracy was similar across cue locations (range: 67% to 77%, all p < 0.001, sign-rank) (Fig 2D).

### Stability of cue information in firing rates

The persistence of cue information in the averaged classification accuracy during the memory period, however robust, may nonetheless belie memory dynamics occurring on single trials. In particular, any lack of persistence on single trials could be obscured in the trial-averaged accuracy. To investigate this, we adapted techniques recently employed to study value coding (Rich & Wallis, 2016) to examine the single-trial dynamics of cue information during working memory. Specifically, we analyzed the confidence of the classifier described above, which provides a time-resolved index of the amount of cue information in population spiking during each trial (Supplementary Materials). In both monkeys, classifier confidence correlated with reaction time on correct trials (Fig. S2). Clearly, if indeed cue information persists on single trials, then confidence values should remain stably above chance (0.5) throughout the memory period.

On the contrary, we found that confidence failed to persist through the memory period on single trials. Instead, lapses in classifier confidence were evident throughout the memory period and across trials within each recording session (Fig. 3A, S3). At the start of each trial, confidence was consistently high during the visual response to the cue. However, following the disappearance of the cue, confidence often returned to chance multiple times during the memory period. During single trials, periods of high confidence were interrupted by sharp transitions to low confidence (Fig. 3B). Lapses in confidence were not associated with microsaccades (Fig. S4). Furthermore, these transitions between high and low confidence did not appear aligned across trials (Fig. 3A,B, S3).

**Figure 3.**
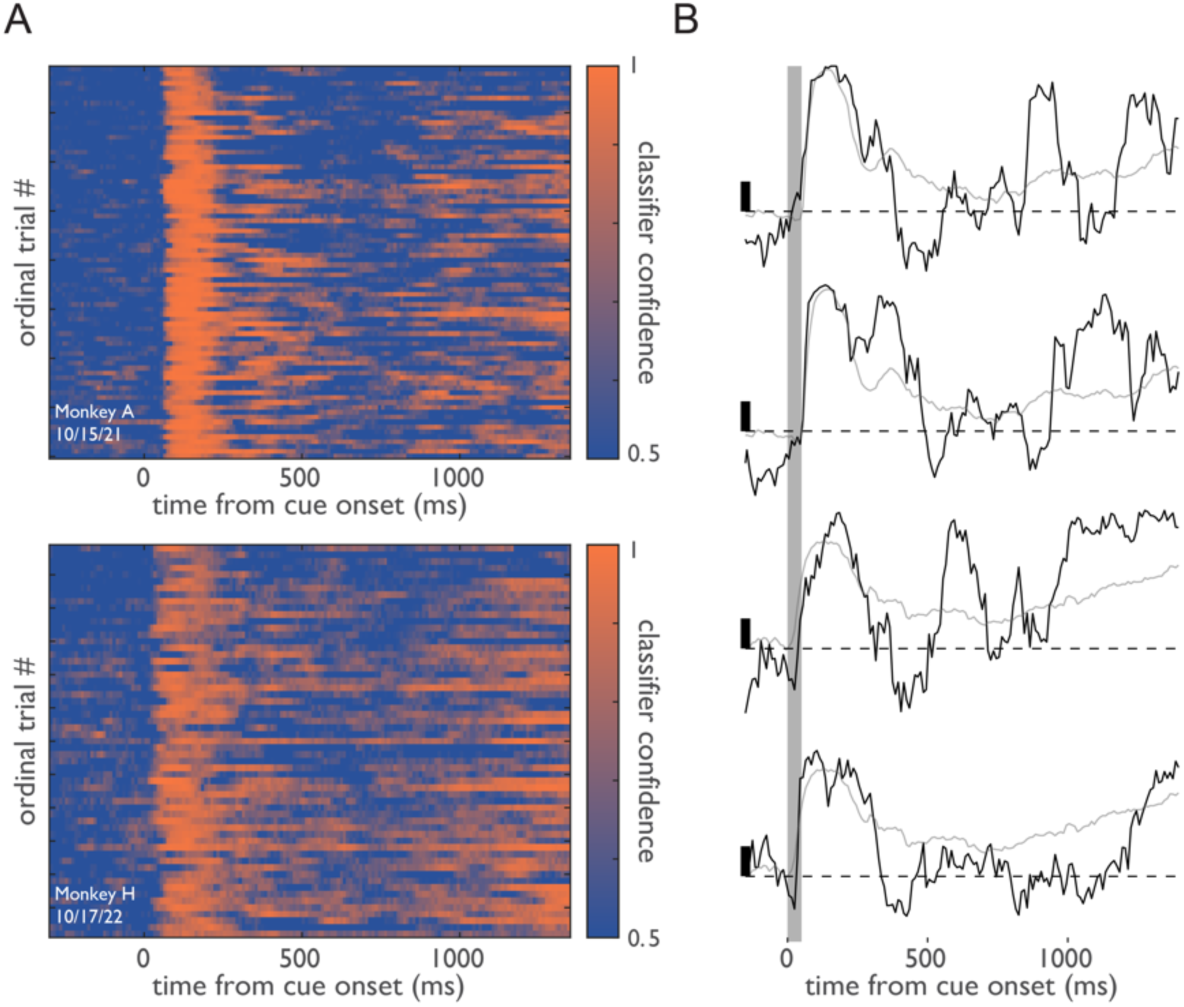
Single-trial dynamics of memory signals in population spiking activity. (A) Single-trial classifier confidence, relative to cue onset, for all trials from the preferred cue condition for two sessions. (B) Confidence (black traces) for four example trials drawn from the two sessions in A. Gray traces show trial-averaged confidence values. Left, black scale bars denote a 0.10 increment in confidence; dashed lines denote y = 0.50.

Given the apparent fluctuations between high and low confidence, we next sought to determine if single-trial confidence was best described by one or two means. Our null hypothesis was that fluctuations in confidence reflected random perturbations around a single mean. We formalized this by fitting a single beta distribution, which is used to model the behavior of random variables on the interval [0, 1], to the histogram of confidence values from each session (Supplementary Materials). The alternative, two-mean, model describes confidence using a mixture of two beta distributions, reflecting two discrete states. Indeed, we found that the two-state model outperformed the single-state model in 15 of 18 recording sessions. This was due to the inability of the single state-model to capture the broad distribution of confidence values (Fig. S5). This asymmetry in model performance was significant across recording sessions (χ^2^(1) = 8, p = 0.005).

### Intermittent rate coding of memoranda is coordinated across the population

Having identified evidence of two discrete states, we next sought to label them on individual trials. We repeated the above classification procedure 50 times, shuffling condition labels on each iteration, to obtain a null distribution of confidence values for each trial (Fig. 4A)(Supplementary Materials). Across the cue and delay epochs, we labeled contiguous time points in which confidence was significantly greater than the null as ‘On’ states, and labeled contiguous nonsignificant time points (p > 0.20) as ‘Off’ states. During each trial, we observed a mean of 2.59 +/- 0.04 (median=3) On states, and a mean of 3.68 +/- 0.02 (median=4) Off states from the cue period until the end of the memory delay (Fig. 4B). Periods in which confidence was significantly below the null (confidently incorrect) were rare (mean=0.14 +/- 0.01 per trial). The mean duration of On states was 197.4 +/- 2.4 (median=160) ms, and the mean duration of Off states was 140.0 +/- 0.6 ms (median=100)(Fig. 4C).

**Figure 4.**
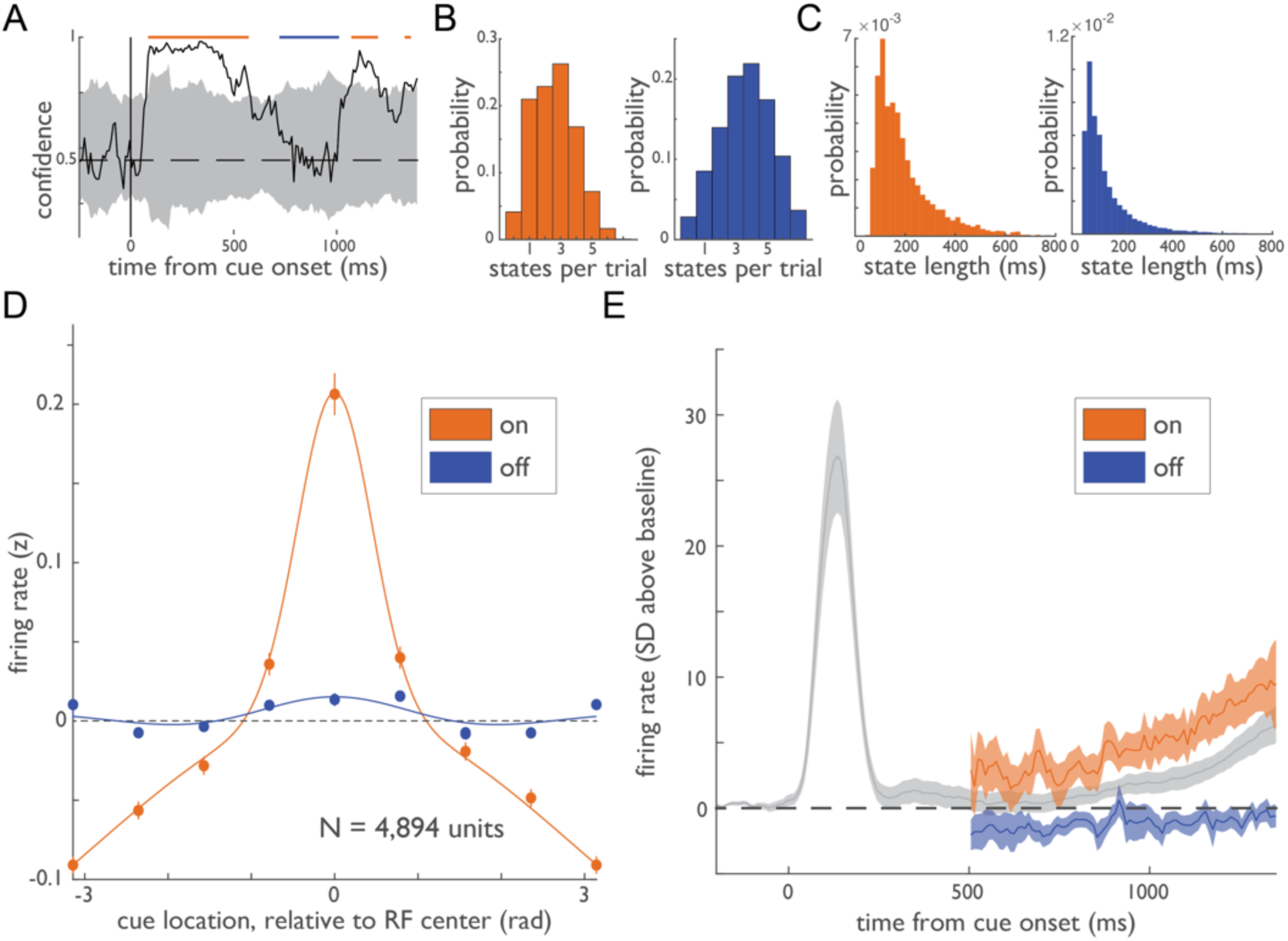
Coordinated changes in memory selectivity and firing rates during On and Off states. (A) Example trial illustrating the labeling of On and Off states. (B) Histogram of number of On (orange) and Off (blue) states per trial and (C) state duration for On and Off states. (D) Memory tuning functions for held-out units during On and Off states. Tuning functions show the mean normalized firing rate during the memory period (z-scored across trials) for held-out units, relative to each unit’s preferred cue location. Error bars (small) denote SEM. (E) Mean normalized population firing rate (SD above baseline) across all sessions for held-out units, relative to cue onset. Averages are plotted for all data points (gray) and also separately for firing rates extracted from On and Off states during the memory period.

Having labeled On and Off states in this way, we next asked how the two states were reflected in the activity of individual neurons during the memory period. To do this, we used half of the neurons recorded during each session to label states as On and Off and then examined the activity of neurons in the remaining (held-out) half of the population from the same sessions. We then repeated this process, switching training and test labels, allowing unbiased analysis of all 4,894 units (Supplementary materials). Indeed, we found that activity differed dramatically between On and Off states in two ways. First, spatial tuning during the memory period, a hallmark of memory delay activity (Rao et al., 1999, 2000), depended heavily on state. During On states, held-out neurons were strongly tuned to the location of the remembered cue (Fig. 4D). In contrast, during Off states, spatial tuning was virtually eliminated (Fig. 4D). Accordingly, there was a significant interaction between cue location and state (On/Off) on firing rates (2-way repeated measures ANOVA, p < 0.001). Whereas cue location explained an average of 9.3% of the variance in firing rate during On states, it explained only 0.8% during Off states, a nearly 12-fold decrease. Thus, Off states were associated with a pronounced loss of spatial tuning at the level of individual neurons.

Second, average firing rates during the memory period also depended on state. During On states, held-out neurons exhibited firing rates above baseline during the memory period (Fig. 4E)(p = 0.002, sign-rank). In contrast, during Off states, firing rates were statistically indistinguishable from baseline (p = 0.145) and significantly lower than On states (p = 0.003). Thus, Off states were not only associated with a loss of spatial selectivity but also a collapse of firing rates to baseline levels. Together, these results reveal how transitions between On and Off states, derived from confidence values, reflect changes in basic firing-rate properties of individual neurons. Furthermore, these observations reveal that transitions between On and Off states during the memory period were coordinated across neurons within the local population.

### Cue-specific neuronal ensembles during the memory period

The fact that cue information carried by neuronal firing rates is periodically lost during the memory period suggests that persistent activity may not be sufficient to support working memory. Therefore, we next looked for evidence in favor of synaptic models. These models propose that, rather than persistent activity, working memory is instead represented by cue-specific networks of neurons (Lundqvist et al., 2011; Mongillo et al., 2008; Stokes, 2015)(Fig. 5A). That is, cue-specific neuronal ensembles should be a signature of working memory. Thus, we next looked for cue-specific cell assemblies during the memory period. As in our analyses of firing rate dynamics, we leveraged the Neuropixels recordings to measure functional connections among the very large numbers of simultaneously recorded neuronal pairs (mean = 36,730 +/- 10,885 per session). Specifically, we examined neuronal cross-correlations to assess their dependence on working memory.

**Figure 5.**
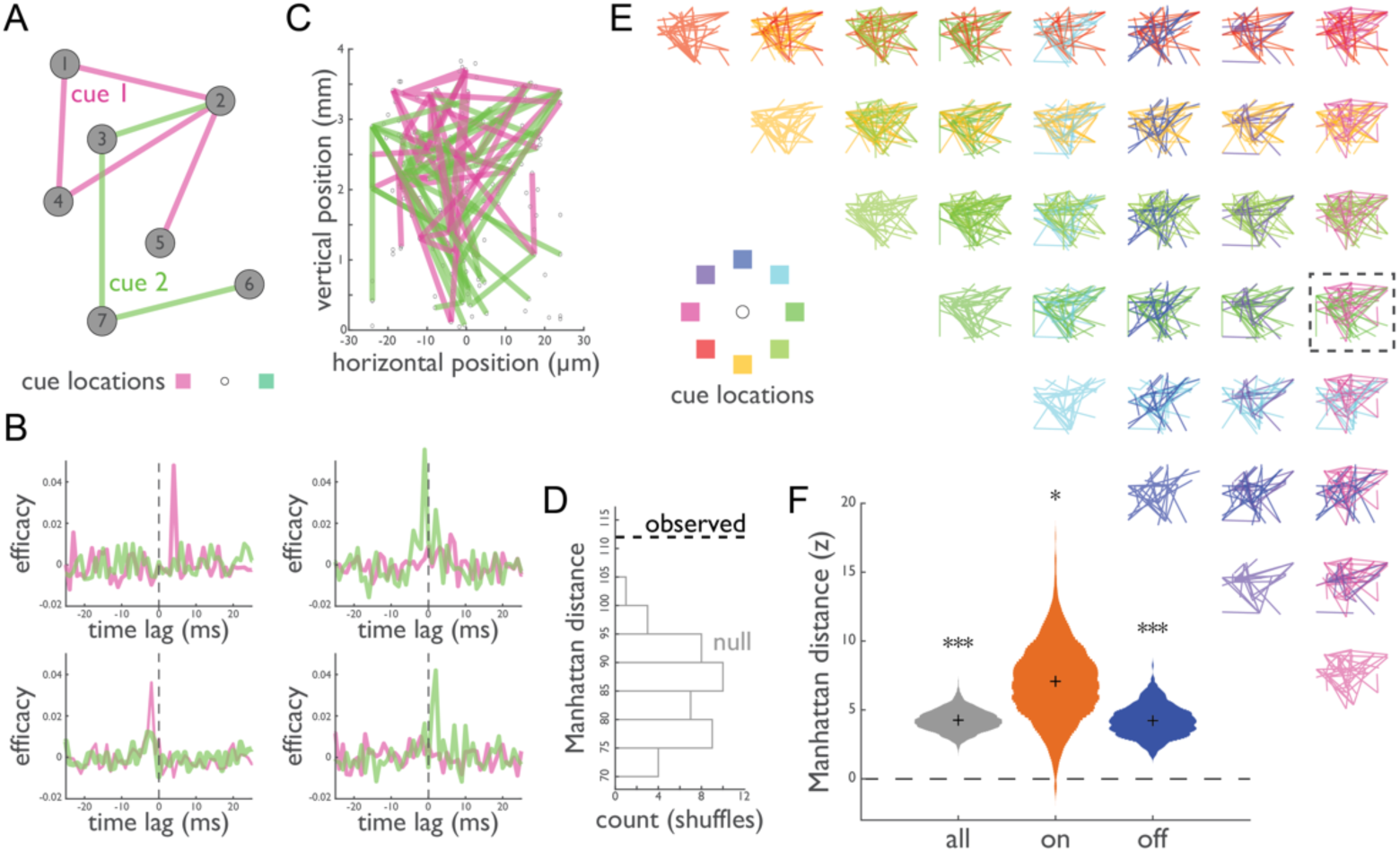
Cue-specific neuronal ensembles during the memory period. (A) Cartoon depicting the synaptic model of working memory. In the absence of cue-selective firing rates, information persists in the cue-specific patterns of potentiated connections (pink and green lines) among neurons (gray nodes). (B) Examples of pairwise CCGs computed during the memory period following two different cues. (C) CCG-derived connectivity maps during the memory period for two different cue locations measured in one session. Lines are drawn between neuronal pairs exhibiting significant CCGs. (D) Differences (dashed line) between the two connectivity maps in C, quantified as the Manhattan distance: the sum of cue-specific connections. Comparison of this metric to a null distribution derived from condition-shuffled data (gray-outlined bars), yielding a z-score (in this case, z = 2.85). (E) Binary comparisons of connectivity maps across all cue locations. (F) Mean normalized Manhattan distance, using data from all sessions during the entire memory period (gray), only during On states (orange), and only during Off states. Violin plots show bootstrap across sessions.

To do this, we computed the cross-correlogram (CCG) between all neuronal pairs in each experimental session and for each cue condition based on activity during the memory period. CCGs were computed and thresholded using established methods (Siegle et al., 2021; Trepka et al., 2022)(Fig. 5b)(Supplementary Materials). We focused on CCGs with non-zero time lags, which are those most consistent with synaptic connections (Ostojic et al., 2009). In our recordings, we observed significant CCGs of this type in 1.52% (+/- 0.25) of neuronal pairs, which totaled 5,868 significant CCGs across all sessions. Next, we compared the pattern of CCGs across cue conditions. Fig. 5c shows an example of a comparison of significant CCGs computed for two cue conditions (0 and 180°) in one recording session. In this example, the two conditions exhibited highly dissimilar ensembles of functionally connected neurons. To quantify the dissimilarity, we counted the number of condition-unique CCGs, yielding the Manhattan distance, which could then be compared to a null distribution derived from condition-shuffled data (Fig 5d). We then repeated this procedure for all possible pairs of conditions in each session (Fig. 5e). Across sessions, the mean Manhattan distance was significantly greater than that predicted by chance (Fig. 5f). Notably, this effect remained significant when confined to a comparison of firing-rate matched conditions (Fig S6b; Supplementary Materials) (p < 0.001, sign-rank). Thus, the ensemble of functionally connected neurons significantly depended on the remembered cue.

If the observed cue-specific ensembles help span the memory period, then they should be evident even when cue information in firing rates is absent. To test this, we repeated the above analysis separately for On and Off states. As expected, the Manhattan distance was significantly greater than chance during On states (Fig. 5f, p = 0.018, sign-rank). Critically, during Off states, when spatial tuning was virtually absent and firing rates transitioned to baseline (Fig.4d,e), the Manhattan distance was also significantly greater than chance (p < 0.001, sign rank). Both On and Off effects remained significant when confined to comparisons of firing-rate matched conditions (Fig S6b; Supplementary Materials) (p = 0.014 and 0.001, respectively; sign-rank). Furthermore, the effect during Off states was indistinguishable from that of On states (p=0.287, sign rank). Thus, even in the absence of persistent memory period activity, memoranda information was reflected in the cue-specific ensembles of functionally connected neurons.

How might spiking and activity-silent coding work together to support memory? Synaptic models of working memory propose that evoked responses to a memory cue potentiate synapses between cue-selective neurons via STP (Fig 6a). During the subsequent memory period, this evoked response relaxes to baseline. However, the cue-specific pattern of potentiated synapses remains. Consequently, cue-specific elevations in firing rate may nonetheless reemerge due to nonspecific fluctuations in extrinsic or intrinsic activity. Thus, across a memory period, multiple transitions between spiking and activity-silent modes (On and Off) may occur (Lundqvist et al., 2011; Mongillo et al., 2008).

**Figure 6.**
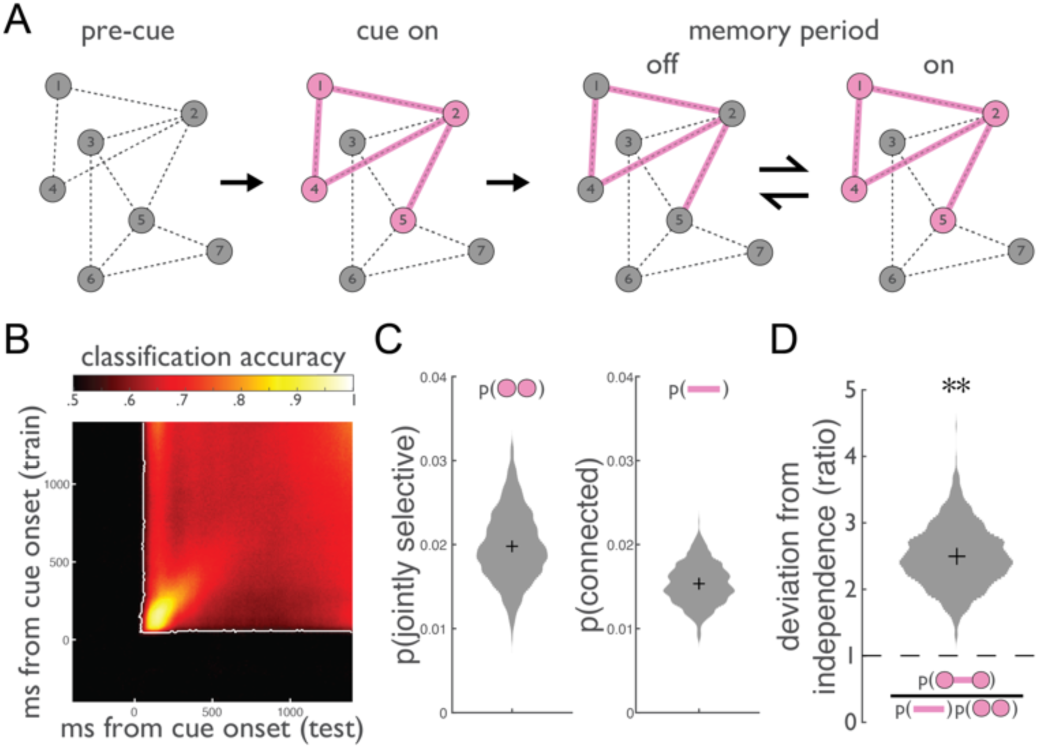
Synaptic model predictions: stability of spike coding and cue-specificity of neuronal ensembles. (A) Cartoon depicting the interplay between spiking and ‘silent’ mechanisms during working memory from synaptic models of working memory. Following the pre-cue period, the memory cue evokes a distinct pattern of activity among a network of neurons (circles) and a distinct pattern of STP, temporarily facilitating connectivity (lines) among cue-associated neurons. During the memory period, even in the absence of persistent spiking activity (Off states), cue information persists in the distinct pattern of connections. During On states, nonspecific drive reignites spiking activity among cue-associated neurons. Paired arrows denote stochastic transitions between Off and On states. (B) Mean classifier accuracy plotted over time points on which the classifier was trained and tested. The block structure indicates good generalization across time. White line: boundary of above-chance classification (corrected for multiple comparisons). (C) Left: probability that a pair of neurons both responded preferentially to the same cue during the evoked response (0-400 ms post-cue). Right: probability that a pair of neurons both exhibited a significant CCG to a particular cue during the memory period. (D) Proportion of pairs that were both jointly selective and connected, divided by the proportion expected. Violin plots indicate bootstrap across sessions.

This account makes two testable predictions. First, it predicts a stable memory code during periods of spiking. That is, despite the interruption of Off states, the same pattern of cue-selective spiking activity should support memory throughout the delay period. Indeed, similar to previous studies (Spaak et al., 2017), we found that the performance of classifiers trained on cue-evoked responses successfully generalized across the entire memory delay period (Fig 6B). Second, there should be a correspondence between spiking responses evoked by the cue and functional connectivity observed during the memory period. Thus, two neurons exhibiting evoked responses to a given cue should tend to be functionally connected during memory. In our data, 2.0% (+/- 0.4) of neuronal pairs responded preferentially to a given cue. Additionally, 1.5 (+/- 0.3)% of neuronal pairs exhibited significant CCGs to a given cue (Fig. 6c). If these proportions are independent, their conjunction should occur at a rate equal to their product. On the contrary, we found pairs displaying spiking responses and significant CCGs to the same cue were observed 2.5 times more than expected (Fig. 6d)(p = 0.003, sign-rank). Thus, neurons that responded jointly during the evoked response were more likely to be functionally connected during the memory period, consistent with synaptic models.

## Discussion

These results elucidate the role of persistent and activity-silent mechanisms in spanning memory periods. By measuring the spiking of large, local populations of prefrontal neurons and resolving the dynamics of mnemonic information on single trials, we found that this information does not persist through the memory period. Instead, cue-specific spiking activity was intermittent, and was characterized by stochastically occurring, discrete transitions between robust coding of memoranda (On states) and complete lapses in such coding (Off states). Notably, these transitions were coordinated across large, local populations of neurons. Complementary to the spike-rate based coding, patterns of functional connectivity also carried information about the remembered cue, even during Off states. These patterns in functional connectivity are consistent with synaptic models of working memory in which cue-specific patterns of potentiated synapses facilitate the reemergence of memory information in spike rates following silent epochs (Lundqvist et al., 2011; Mongillo et al., 2008; Stokes, 2015).

Our observations describe the dynamics of local populations of neurons in the primate dorsolateral prefrontal cortex. Future work should explore these dynamics in other structures known to exhibit memory-delay activity, e.g., parietal cortex (Snyder et al., 1997), and on broader spatial scales, including the possible propagation of on states across the cortical surface (Shi et al., 2022) or distributed representations supporting memory (Li et al., 2016; Schmitt et al., 2017). Nevertheless, two facts suggest that the regions targeted by the present study play a key role in working memory. First, focal inactivation of dorsolateral prefrontal cortex has long been known to impair performance in spatial working memory tasks (Chafee & Goldman-Rakic, 2000; Dias & Segraves, 1999; Sommer & Tehovnik, 1997), including selective inactivation of memory-delay activity (Acker et al., 2016). Second, in the context of our study, single trial confidence at the onset of the behavioral response phase predicted behavior.

Similar, large-scale electrophysiological approaches should be used to assess the relative roles of spiking and synaptic mechanisms across a range of working memory tasks. For example, such roles may be rather different for spatial and object-based working memory, given the apparent differences between the two in the robustness of memory-delay spiking activity across brain areas (Christophel et al., 2017). Furthermore, theoretical work suggests that the relative contribution of these two mechanisms may be related to the amount of manipulation of remembered information demanded by a task (Masse et al., 2019). The spatial tasks employed here demand a relatively straightforward sensory-to-motor transformation, and so could rely more on synaptic mechanisms. Empirically verifying how task demands sculpt memory representations will lead to a richer understanding of the mechanistic basis of working memory.

Finally, it is interesting to note that even within a single experimental session and task condition, the relative proportion of On and Off states showed a fair degree of heterogeneity across trials. Recent electrophysiological studies provide evidence of coordinated fluctuations in local neuronal activity in alert animals (Harris & Thiele, 2011). Furthermore, these coordinated fluctuations appear related to moment-to-moment changes in global arousal states and also predict psychophysical performance in nonhuman primates (Davis et al., 2020; Engel et al., 2016). Among the many possible mechanisms that underlie coordinated fluctuations in neuronal activity include neuromodulatory inputs (Noudoost & Moore, 2011). Dynamics in the local tone of neuromodulators may be sufficient to induce transitions in local cortical states (Harris & Thiele, 2011). Within dorsolateral prefrontal cortex, dopaminergic tone is known to play a key role in the maintenance of memory-delay activity (Sawaguchi & Goldman-Rakic, 1991; Vijayraghavan et al., 2007). Thus, examining the contribution of dopaminergic tone to the variability in the dynamics of memoranda coding could prove illuminating.

## Acknowledgements

The authors would like to thank Danielle Lopes and Stephen Cital for veterinary assistance, Mehmet Solyali and Bob Schneeveis for machining assistance, and Tatiana Engel, Shaul Druckmann, Alireza Soltani, and Xiaomo Chen for helpful comments. This work was supported by NIH grants EY014924 and NS11662302, and a Ben Barres Professorship to T.M..

## Data and code availability

The data and custom code supporting this study will be made available online upon final publication.

## Author Contributions

Conception: MP, TM. Task design: DJ, TM. Animal Training: DJ. Surgical Procedures: TM, DJ. Data collection: MP, DJ. Analysis: MP, JO. Software Tools: SZ, ET. Supervision: TM. Funding Acquisition: TM. Writing – first draft: MP, TM. Editing and revision – all authors.

## Supplementary Materials

### Supplementary Methods

#### Subjects

Two adult male rhesus monkeys (*Macaca mulatta*) participated in the experiment. Monkey A and H weighted 11 and 14 kg, respectively. All surgical and experimental procedures were approved by the Stanford University Institutional Animal Care and Use Committee and were in accordance with the policies and procedures of the National Institutes of Health.

#### Behavioral task

Stimuli were presented on a VIEWPixx3D monitor positioned at a viewing distance of 60 cm using Psychtoolbox and MATLAB (MathWorks). Eye position was monitored at 1kHz using an Eyelink 1000 eye-tracking system (SR Research). On each trial, the animals were presented with a cue at one of 8 possible locations and reported this location after a brief memory delay to receive fluid reward. Cues were square frames (green for Monkey A, black for Monkey H) measuring 1 degree of visual angle on a side and presented at 5-7 degrees of eccentricity (depending on the session).

Monkeys initiated behavioral trials by fixating a central fixation spot presented on a uniform gray background. After the monkeys maintained fixation for 600-800 ms (randomly selected on each trial), a cue appeared for 50 ms at one of 8 possible locations separated by 45 degrees around fixation. Cue presentation was followed by a delay period that varied randomly from 1400-1600 ms. After the delay period, the fixation spot disappeared, and the animal was presented with one of two possible response screens. On match-to-sample (MTS) trials, two targets appeared (filled blue circles, radius 1 DVA), one at the previously cued location, and the other at one of the 7 remaining non-cued locations. On memory-guided saccade (MGS) trials, no targets appeared. In either case, the animals received a juice reward for making an eye movement to within 5 degrees of visual angle (DVA) of the previously cued location and then maintaining fixation for 200 ms. MTS and MGS trials were randomly interleaved such that the animals could not predict the trial type. The animals had to maintain their gaze within 3 DVA (monkey A) or 2 DVA (monkey B) from fixation throughout the trial until the response stage. The intertrial interval was 300-600 ms after each correct response. Failures to acquire fixation, fixation breaks, and incorrect responses were not rewarded and were followed by a 2,000 ms intertrial interval.

#### Surgical Procedures and Recordings

Monkeys were implanted with a titanium headpost to immobilize the head and with a titanium chamber to provide access to the brain (see Armstrong et al., 2009 for full details). In a previous study (Jonikaitis et al., 2023), we identified the FEF based on its neurophysiological characteristics and the ability to evoke saccades with electrical stimulation. Here, we recorded from FEF and anterior sites (Broadmann areas 9/46) using primate neuropixels probes (Trautmann et al., 2023). We pierced the dura using a screw-driven 21 gauge pointed cannula and lowered the probe through this cannula using a combination of custom 3D printed grids and motorized drives (NAN instruments). Recordings were allowed to settle for ∼30 minutes prior to the start of the experiment to mitigate drift. We configured probes to recorded from 384 active channels in a contiguous block, allowing dense sampling of neuronal activity along a 3.84 mm span.

Neuronal waveforms identified using automated spike-sorting routines (Kilosort3, Pachitariu et al., 2023). Neuropixels filter and digitize activity at the headstage separately for the action potential (300 Hz high-pass filter, 30 kHz sampling frequency) and local field potential (1 kHz low-pass filter, 2.5 kHz sampling frequency) bands. Activity was monitored during experimental sessions and saved to disk using SpikeGLX (https://billkarsh.github.io/SpikeGLX/).

#### Data Preprocessing

Spiking in the action-potential band activity was identified and sorted offline using Kilosort3 (Pachitariu et al., 2023). As we were interested in population-level coding of memory, we analyzed both putative single- and multi-unit clusters identified by Kilosort. Spike times were aligned to a digital trigger on each trial indicating cue onset and corrected for a lag in stimulus presentation estimated offline using photodiode measurements from the stimulus display and the timing of the cue-evoked response. Neurons that fired fewer than 1000 spikes in the ∼3 hour experimental sessions were excluded from further analyses. Spikes times were converted into smoothed firing rates (sampling interval, 10 ms) by representing each spiking event as a delta function and convolving this time series with a 100-ms boxcar. For cross-correlogram analyses, unsmoothed spikes times were binned with a width and timestep of 1 ms. Incorrect trials were rare (Fig. 1b) and were excluded from subsequent analysis.

#### Functional subtyping

To determine the functional subtype of units (Bruce & Goldberg, 1985)(Fig. S1), we analyzed firing rates during three time epochs: visual (0 to 400 ms post-cue onset), memory (500-1400 ms post-cue onset) and motor (100-300 ms post-fixation offset). A unit was labeled as being selective during a given epoch if firing rates during that epoch were significantly modulated by cue location (1-way ANOVA, p < 0.05 criterion). Units were then sorted into functional subclasses based on the set epochs during which each unit was selective.

#### Classification of cue location

Firing rate estimates for each unit and timepoint relative to cue onset were z-scored across trials prior to the classification. We used linear classifiers to quantify the amount of information about the location of the cue in populations of simultaneously recorded units. We held out each trial for test one by one, training a logistic regression classifier (as implemented by fitclinear.m in MATLAB) to predict the cue location using the population vector of firing rates. Specifically, classifiers were trained to discriminate the same cue location as the test trial from the opposite cue location using the applicable subset of trials from the training set. Data were subsampled during training to equalize trial counts for the two conditions. A unique classifier was trained and tested for each timepoint relative to cue onset. ‘Classification accuracy’ reflects the proportion of correctly classified test trials (Fig. 2c). ‘Classifier confidence’ is the non-thresholded value of the logistic function corresponding to the probability assigned by the classifier to the correct label at test (Fig. 3).

Cross-temporal classification (Fig. 6b) was similar, except that we employed a split-half approach where the classifiers for each timepoint were trained on half of the available population of trials and tested (cross-temporally) using the other half.

#### Mixture modeling of confidence

We used a mixture modeling approach to test whether confidence during the memory period (500 to 1400 ms post-cue) was best described as drawn from a 1- or a 2-state distribution (Fig. S5). To do this, for each session and cue location, we modeled the PDF of confidence values during the memory period as either a single beta distribution:

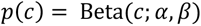

or as a mixture of two beta distributions:

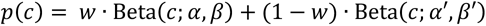

where *c* is confidence, α, β, α’ and β’ parameterize the beta distribution(s), and *w* is the mixing coefficient. The best fitting parameters of each model were identified by maximum likelihood estimation using gradient descent in MATLAB. We used 4-fold cross validation on the population of trials to assess the likelihood of each model on held-out test data, and then normalized by the number of trials and changed the log likelihood to base 2 to yield the cross-validated score of each model in terms of *bits per trial*. Finally, we subtracted these two model scores and averaged across conditions to yield the difference in model performance for each session.

Similar results for our one-vs-two state model comparison were obtained when using gaussian mixture or hidden Markov models. We settled on a beta mixture model approach because the beta distribution was most appropriate for data distributed on the interval [0, 1] and because state transitions were non-Markov (e.g., due to increased likelihood of On states at the end of the delay).

#### Analysis of microsaccades

The horizontal and vertical eye position records were convolved with a Gaussian kernel (σ = 4.75 ms) to suppress noise before taking first derivatives, yielding the eye velocity along each dimension. We then took the root sum of squares of the horizontal and vertical velocities to obtain eye speed. We flagged peaks in this timeseries with a minimum peak height of 10 deg/second and a minimum interpeak distance of 50 ms as microsaccades (Bair & O’Keefe, 1998, Fig. S4), which were confirmed via visual inspection of the data.

#### Labeling of on and off states

To identify On and Off states (Fig. 4a), we repeated the cue classification analysis described above 50 times, randomly shuffling the labels of the training set for each test trial. This yielded, for each trial, a null distribution of 50 confidence timeseries (Fig 4a). We then z-scored each timepoint of the true confidence time series by the mean and standard deviation of this null distribution. Individually significant (>1.96) z-values were cluster-corrected for multiple comparisons over time (Maris & Oostenveld, 2007). In brief, we compared the sum of contiguous individually significant z-values with that expected by chance (randomization test). Clusters with a mass greater than the 95% percentile of the null were labeled ‘On’ states. Contiguous z-values falling below a conservative (p > 0.20) threshold for at least 5 consecutive timepoints were labeled ‘Off’ states.

#### Tuning curves

To test if On and Off states reflected coordinated changes in tuning across the neural population, we used a split-half approach. First, firing rate estimates for each unit and timepoint relative to cue onset were z-scored across trials. Then, for each session, we randomly divided the population of units in half. We used one half of the units to identify On and Off states, as described above. Next, for each unit in the held-out population, we computed the mean firing rate during the memory period for each cue location separately for On and Off states, averaging across relevant timepoints and across trials. This yielded, for each unit, two 8-element vectors – the On and Off tuning functions. To align tuning functions across units, the preferred cue location for each was identified as condition in which the sum of the On and Off functions was greatest and assigned an arbitrary value of zero degrees. Aligning tuning curves to the maximum-valued ‘preferred’ cue in this way will necessarily produce a peak at zero degrees in the average tuning function, even in the absence of true tuning. To correct for this, for each unit we also computed null On and Off tuning functions by first shuffling cue labels across trials, aligned these to the ‘preferred’ cue, and subtracted these from the true On and Off tuning functions (Fig 3d).

For display purposes, we fit the average On and Off tuning functions with a difference of gaussians using gradient descent in MATLAB. Difference-of-gaussians are useful for describing tuning curves that display surround suppression (Sceniak et al., 2001).

#### Population firing rates

To described how population firing rates evolved over the course of the trial, we averaged firing rates across all units recorded in the same session and across all trials for the preferred cue location (greatest mean classification confidence during the memory period), yielding a single time series for each session. We then normalized this time series by the mean and standard deviation of a 400 ms baseline period (-400 to 0 ms relative to cue onset), yielding a metric of population spiking in units of standard deviations above baseline (Fig 4e, gray traces). We repeated this analysis for the memory period, this time only including datapoints labeled On or Off (Fig 4e, orange and blue traces).

#### Cross-correlogram analysis

To characterize functional connectivity among units, we computed the crosscorrelation between spike trains of all pairs of simultaneously recorded neurons with mean firing rates greater than 1 1 Hz. CCGs were computed separately for each cue location. Following previous studies (e.g. Trepka et al., 2022), to mitigate the firing rate effects, we normalized the crosscorrelation for each pair of neurons by the geometric mean of their firing rates. The cross-correlogram (CCG) for a pair of neurons (*j, k*) in condition *c* was therefore:

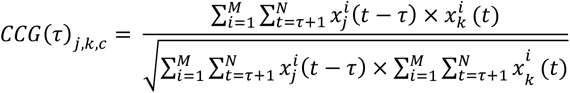

where *M* is the number of trials collected for cue location *c*, *N* is the number of time bins within a trial, τ is the time lag between the two spike trains, and *x_k_^i^*(*k*) is one if neuron *j* fired in time bin *t* of trial *i* and zero otherwise.

To correct for correlation due to stimulus locking or slow fluctuations in population response, we subtracted a jittered cross-correlogram from the original cross-correlogram. This jittered cross-correlogram reflects the expected value of the cross-correlogram computed from all possible jitters of each spike train within a given jitter window (Harrison & Geman, 2009; Smith & Kohn, 2008). The jittered spike train preserves both the PSTH of the original spike train across trials and the spike count in the jitter window within each trial. As a result, jitter correction removes the correlation between PSTHs (stimulus-locking) and correlations on timescales longer than the jitter window (slow population correlations). We chose a 25-ms jitter window, following previous work (Jia et al., 2013; Siegle et al., 2021; Trepka et al., 2022; Zandvakili & Kohn, 2015).

We classified a CCG as significant if the peak of the jitter-corrected CCG occurred within 10 ms of zero and was more than seven standard deviations above the mean of a high-lag baseline period (100 > |τ| > 50, Siegle et al., 2021). Zero-lag CCGs were excluded from the analyses reported here, although including them yielded statistically indistinguishable results.

All CCGs were estimated using spike trains during the memory period (500 to 1400 ms post-cue) to avoid the influence of visually-evoked responses. CCG analyses specific to On and Off states (Fig 5f) were computed by first setting *x(t)* to zero for all timepoints not identified as ‘On’ or ‘Off’ (respectively), and then repeating the analysis described above.

#### Manhattan distance

To determine if patterns of functional connectivity differed according to the contents of memory, we compared the graphs of significant CCGs across cue locations in a pairwise manner (Fig. 5c-f). For each session and cue location, we represented the results of our CCG analyses as a graph in which nodes were units. The edge (connection) between each pair of units was assigned a weight of one if the pair had a significant CCG and zero otherwise. Then, for each possible pair of cue locations, we computed the Manhattan distance, the number edges with a weight that differed across the two graphs. Finally, we averaged this metric across all 28 possible pairs of conditions, yielding one summary statistic per session.

To normalize this mean Manhattan distance for comparison across sessions, we shuffled the cue location labels across trials and repeated the entire analysis pipeline 50 times (25 for analyses specific to On and Off states), from CCG estimation through Manhattan distance calculation. We then z-scored the mean Manhattan distance for each session by this null distribution and compared these z-scores to zero (Fig. 5f).

Note that CCGs among both single- and multi-units have been widely used as a measure of functional connectivity (e.g., deCharms & Merzenich, 1996; Eckhorn et al., 1988; Engel et al., 1990; Gray et al., 1992; Gray & Singer, 1989; Luczak et al., 2015; Tanaka et al., 2014). Indeed, CCGs based on multi-unit activity may be more sensitive at detecting correlations in spiking than similar analyses of single-neuron pairs (Bedenbaugh & Gerstein, 1997; deCharms & Merzenich, 1996; Roy & Alloway, 1999). Nonetheless, the presence of multi-units in our dataset does limit the conclusions that might be drawn about the specific neuronal subtypes involved in the cue-dependent ensembles that we observe, e.g. putative pyramidal vs non-pyramidal neurons.

#### Firing rate matched control

The geometric mean firing rate (gFR) of pairs of units varied significantly across the eight cue locations (1-way ANOVA, p = 0.013)(Fig. 6a). gFRs were statistically indistinguishable, however, across cue locations 1-5 (p = 0.146) and 6-8 (p = 0.593). Therefore, we repeated the analysis of Manhattan distance described above, this time only computing the Manhattan distance among cue locations 1-5 and among 6-8 (Fig. S6b) to yield a firing-rate matched variant of the analysis presented in Fig. 5f.

#### Joint Selectivity

To determine the selectivity of units during the evoked response, we averaged each unit’s cue-locked firing rate over time (from 0 to 400 ms post-cue onset), yielding an nTrials x 1 vector of firing rates. We then performed a one-way ANOVA to evaluate the relationship between cue location and firing rate. If the effect of cue location was significant (p < 0.05), the unit was deemed ‘selective’ to cue location and the location to which it had the greatest mean firing rate was labelled the preferred location. Pairs of units were deemed jointly selective if they were selective for the same cue location.

---

**Fig S1.**
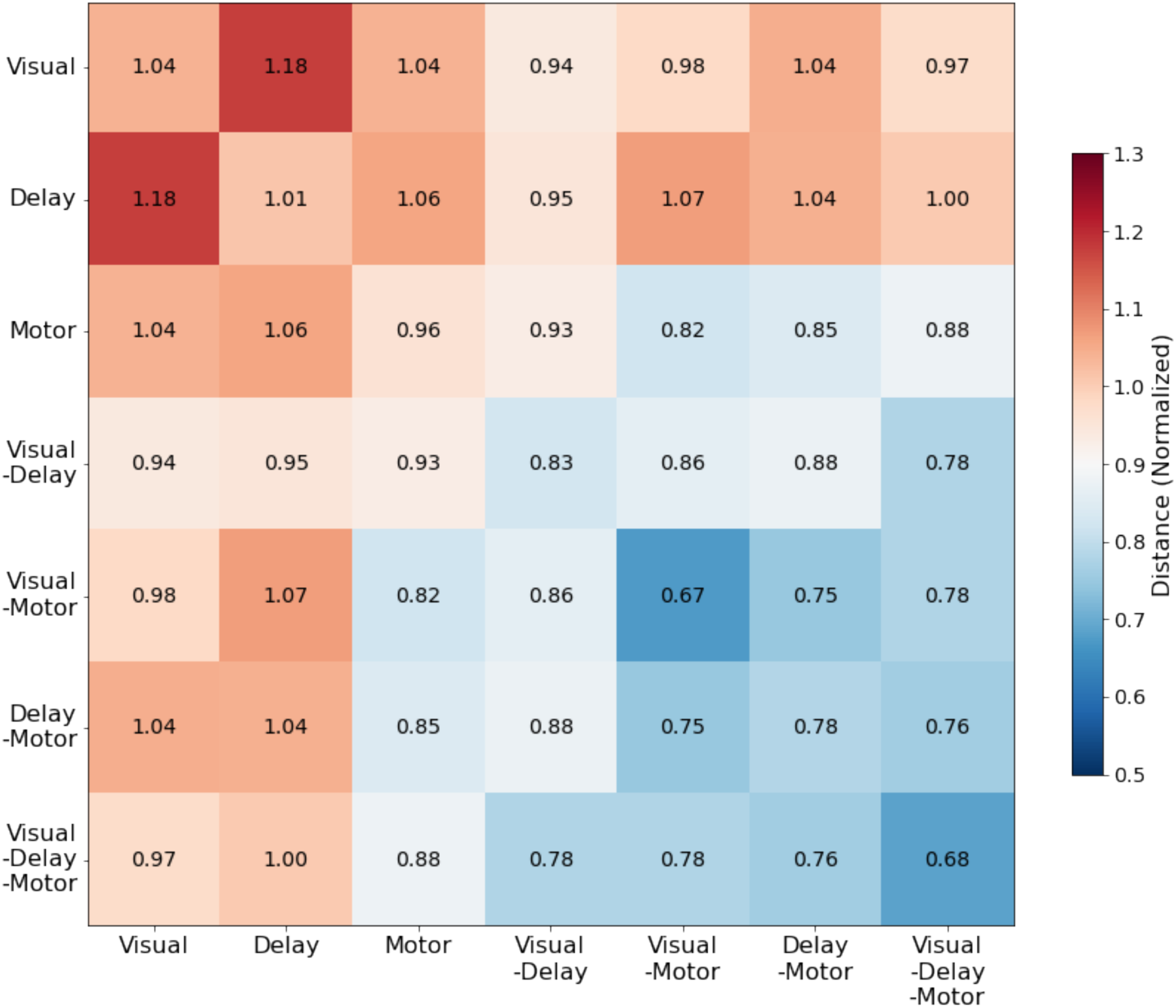
Normalized distance between different functional classes of neurons within lateral prefrontal cortex. Seven functional classes of neurons were defined according to selective activity to three task components: visual, delay, and motor. Neurons were defined as having a given functional property based on the presence of significant selectivity across cue conditions within the visual, delay and motor epochs of the task. The plot shows the mean distance between different classes of neurons; means were calculated and normalized by each session’s total mean distance. Top-left to bottom-right diagonal elements show the mean distance within each functional class.

**Figure S2.**
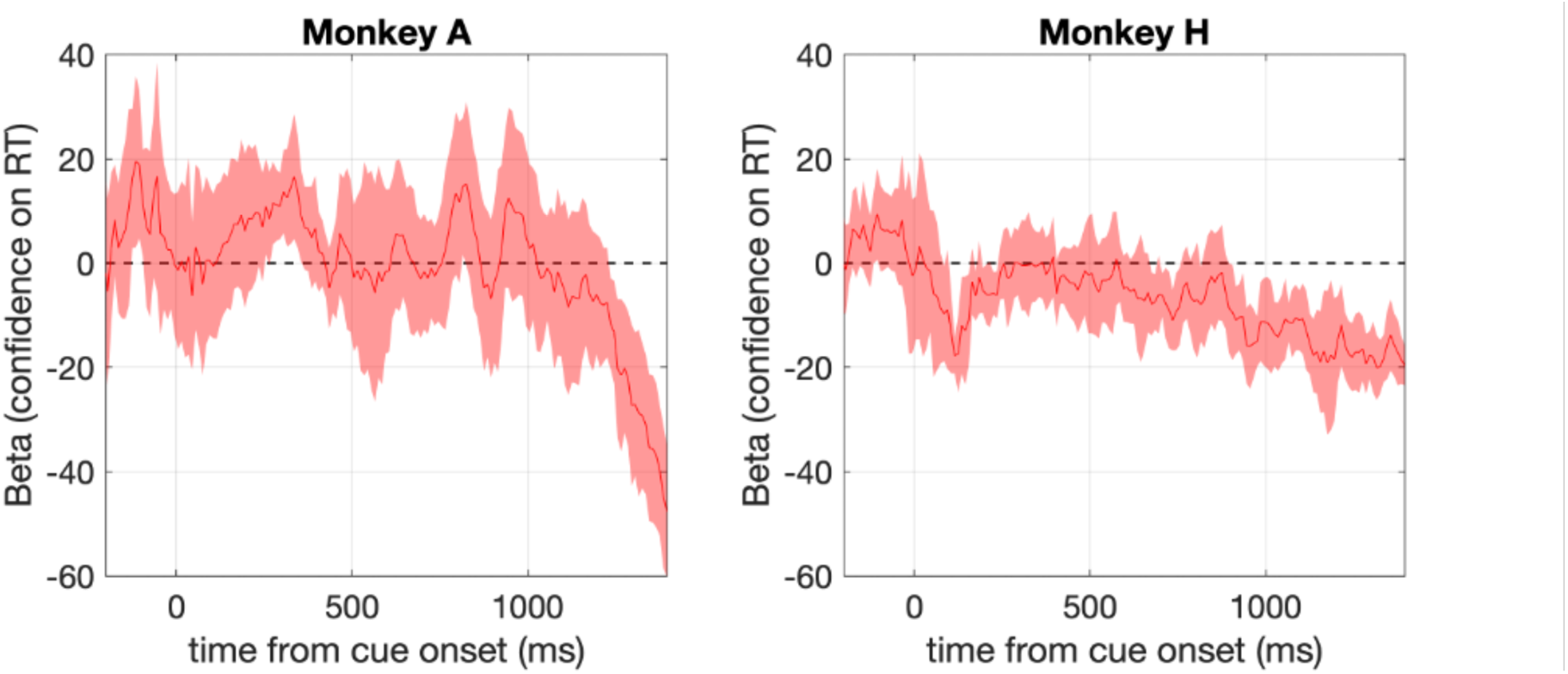
Regression coefficient relating classifier confidence to reaction time in milliseconds (average across N=10 and N = 8 sessions). Cue location and task (MTS/MGS) were included as co-regressors. Shaded area: 95% confidence intervals.

**Figure S3.**
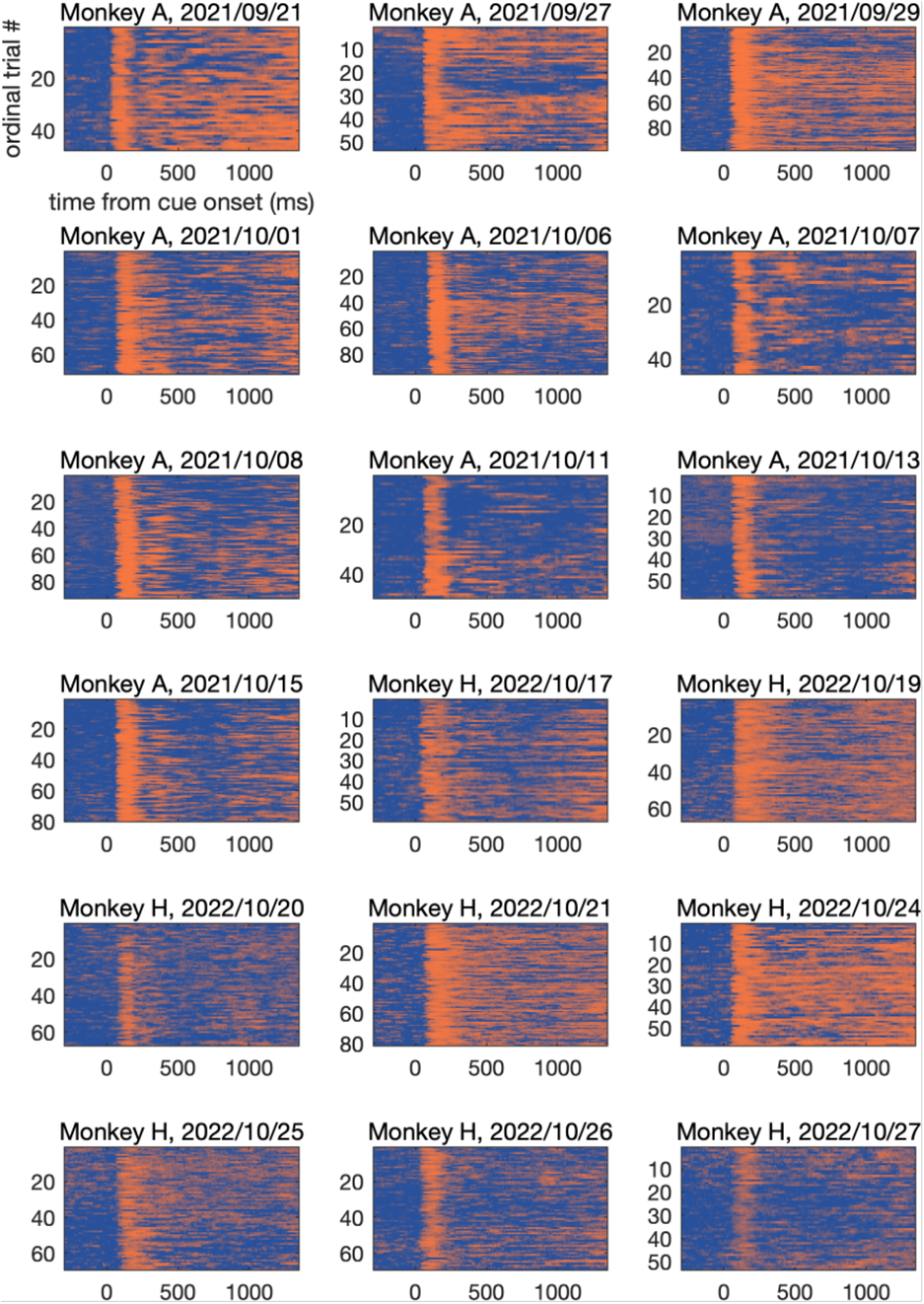
Single-trial classifier confidence, relative to cue onset, for all trials from the most preferred cue condition for each session. Color scale as in Figure 2a.

**Figure S4.**
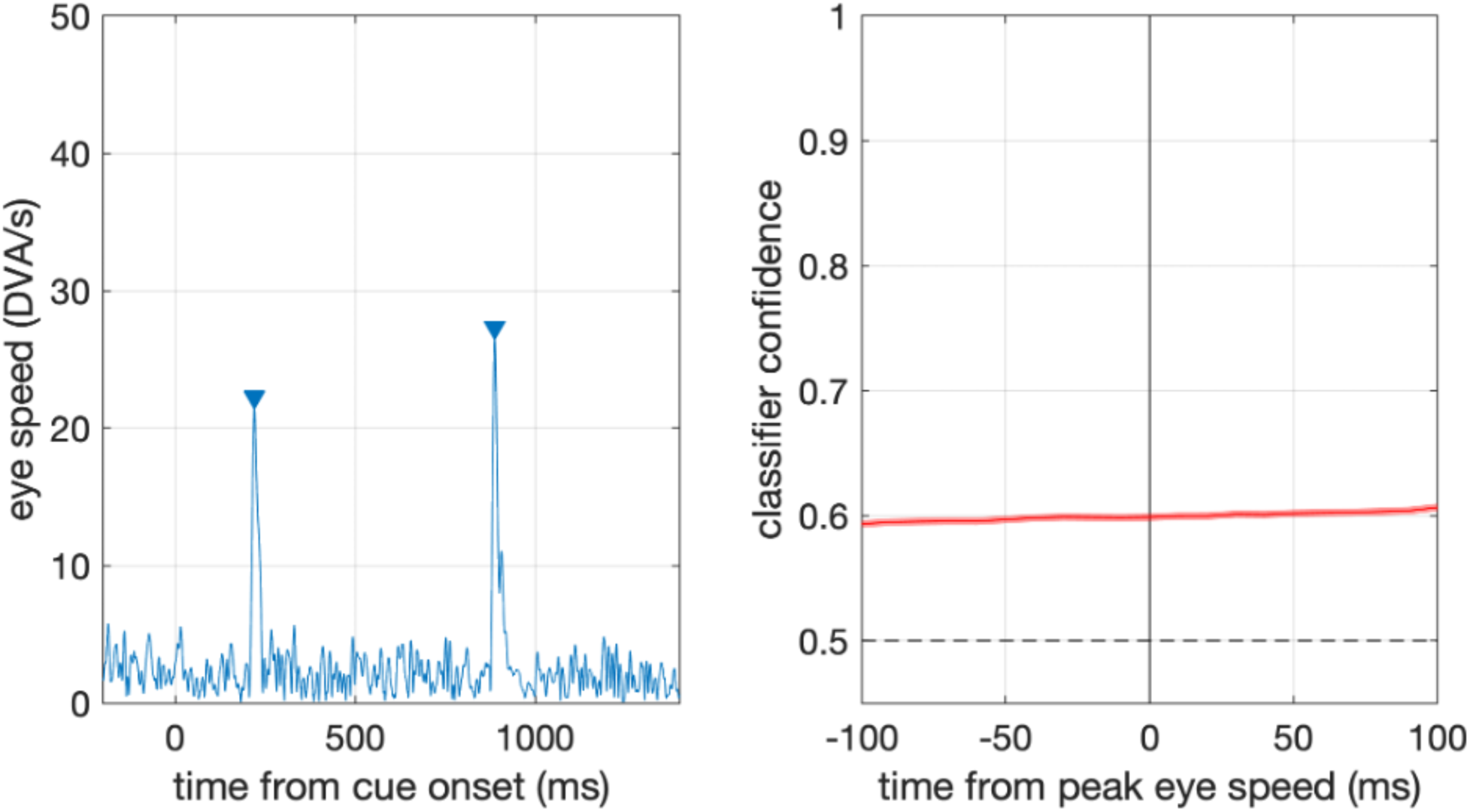
Classifier confidence was not affected by microsaccades. Left: Example trial showing eye speed data and microsaccade identification. Microsaccades were identified as peaks in eye speed >10 DVA/s. Right: Mean classifier confidence during the memory period, locked to microsaccades (average across N = 8,910 microsaccades). Shaded area (small) indicates 95% confidence intervals.

**Figure S5.**
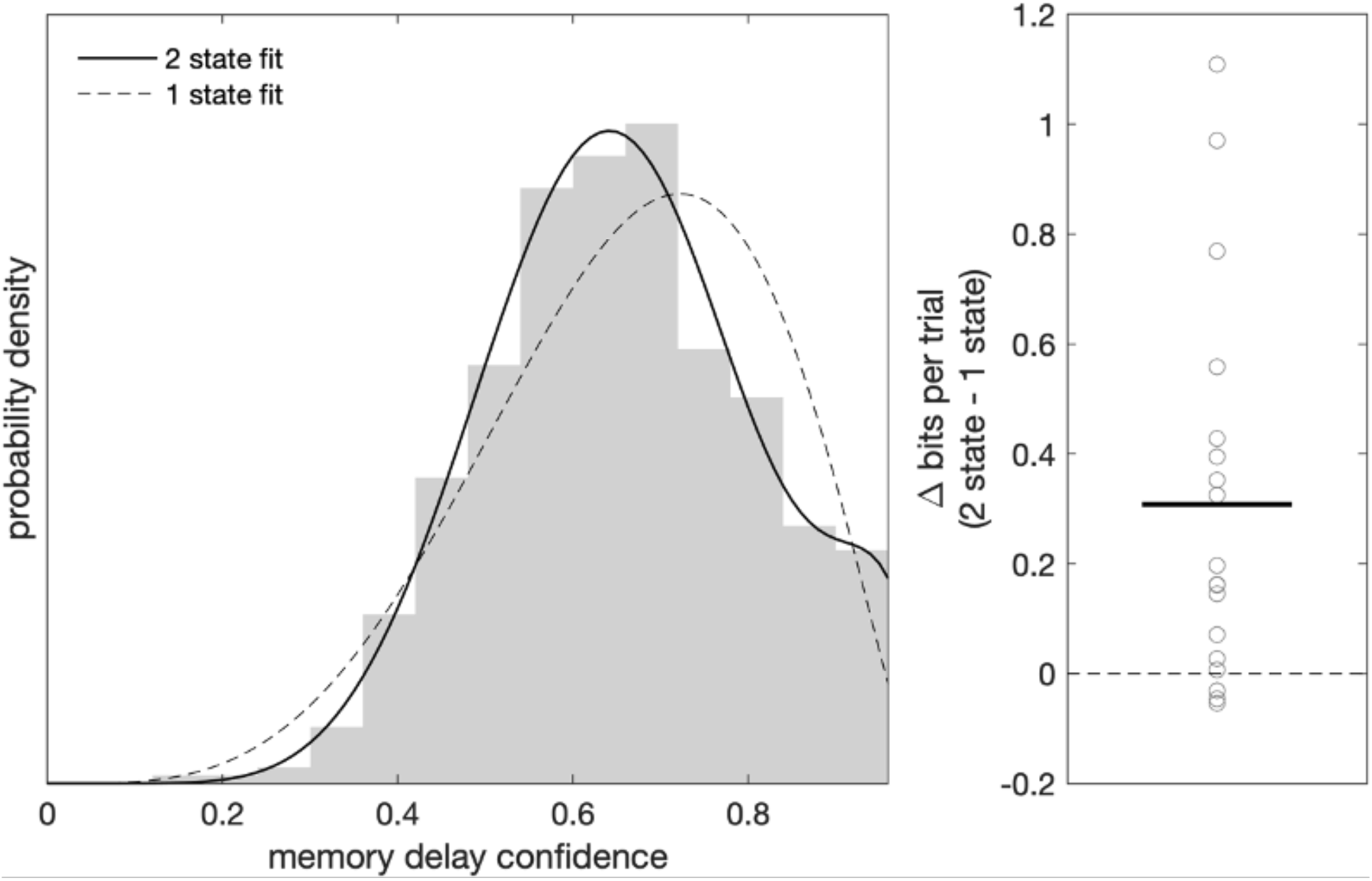
Confidence values during the memory period are better described as draws from a 2-state rather than from a 1-state model. Left: Histogram of confidence values during memory delay time points for the preferred cue condition from one example session. Dashed line shows the best fitting beta distribution. Solid line shows the best fitting mixture of two beta distributions. Right: Cross-validated model comparison results for 2-state vs 1-state fits for all sessions (N=18). Circles show individual session scores; black line shows mean across sessions; dashed line indicates equivalent model performance.

**Figure S6.**
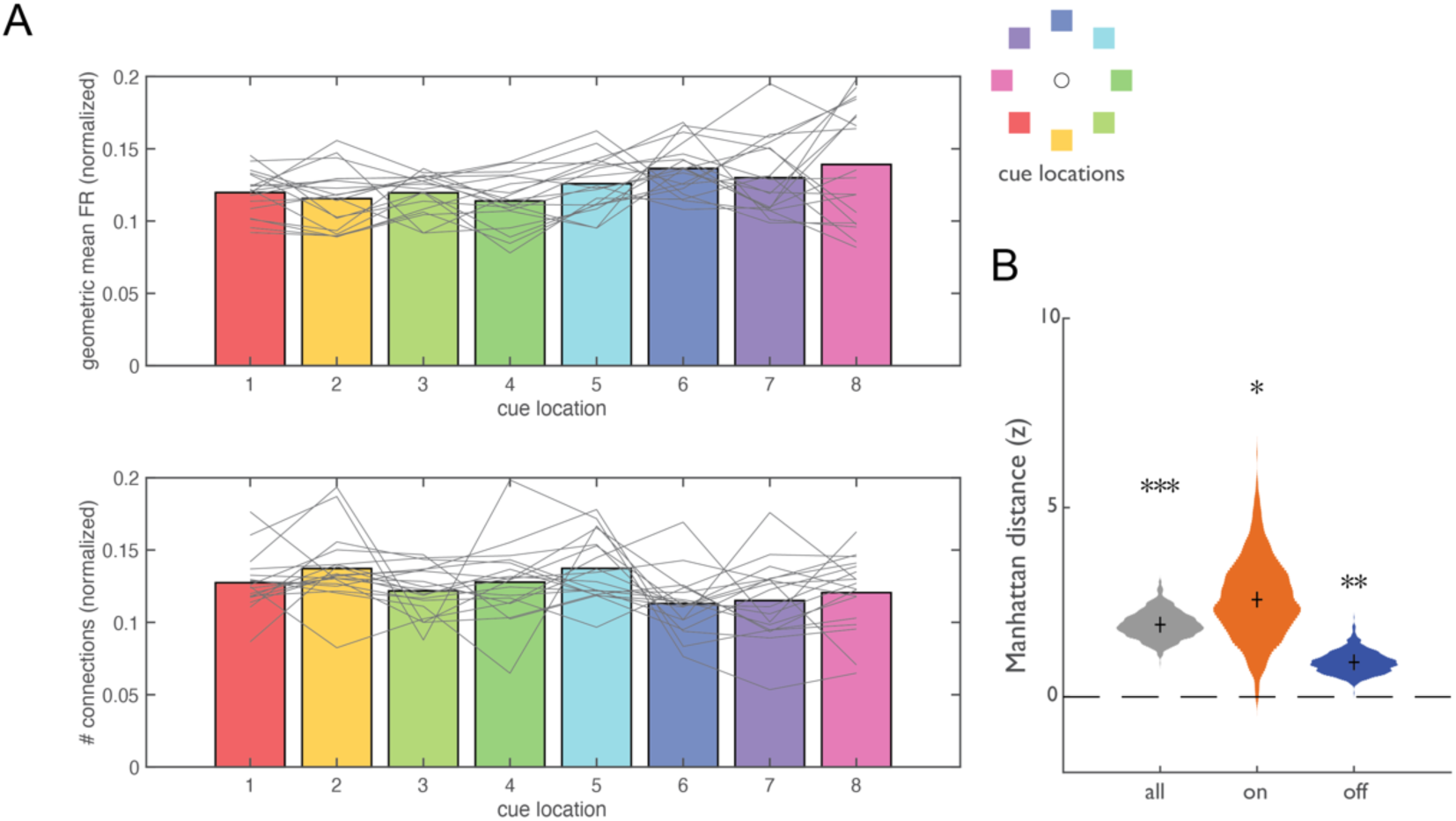
Firing rate matching for CCG analysis. (A) Top: Normalized geometric mean firing rate (gFR) of all pairs of neurons during the memory period. Bottom: Normalized number of neuronal pairs with significant CCGs. Gray lines show individual sessions; bars show mean across sessions; colors indicate cue location (lower inset). (B) Mean normalized Manhattan distances restricted to a comparison among cue conditions (1-5 and 6-8) in which gFR was equal (see Methods). Shown are means of data from all sessions during the entire memory period (gray), only during On states (orange), and only during Off states. Violin plots show bootstrap across sessions.

